# A molecular switch in RCK2 triggers sodium-dependent activation of K_Na_1.1 (KCNT1) potassium channels

**DOI:** 10.1101/2023.09.28.559907

**Authors:** Bethan A. Cole, Antreas C. Kalli, Nadia Pilati, Stephen P. Muench, Jonathan D. Lippiat

## Abstract

The Na^+^-activated K^+^ channel K_Na_1.1, encoded by the *KCNT1* gene, is an important regulator of neuronal excitability. How intracellular Na^+^ ions bind and increase channel activity is not well understood. Analysis of K_Na_1.1 channel structures indicate that there is a large twisting of the βN-αQ loop in the intracellular RCK2 domain between the inactive and Na^+^-activated conformations, with a lysine (K885, human subunit numbering) close enough to form a salt bridge with aspartate (D839) in the Na^+^-activated state. Concurrently, an aspartate (D884) adjacent in the same loop adopts a position within 4 Å of several acidic or polar residues. In carrying out mutagenesis and electrophysiology with human K_Na_1.1, we found alanine substitution of each of these residues resulted in almost negligible currents in the presence of up to 40 mM intracellular Na^+^. The exception was D884A, which resulted in constitutively active channels in both the presence and absence of intracellular Na^+^. Further mutagenesis of this site revealed an amino acid size-dependent effect. Substitutions at this site by an amino acid smaller than aspartate (D884V) also yielded constitutively active K_Na_1.1, D884I had Na^+^- dependence similar to wild-type K_Na_1.1, whilst increasing the side chain size larger than aspartate (D884E or D884F) yielded channels that could not be activated by up to 40 mM intracellular Na^+^. We conclude that Na^+^ binding results in a conformational change that accommodates D884 in the acid-rich pocket, which triggers further conformational changes in the RCK domains and channel activation.

**Statement of Significance:** Sodium-activated potassium channels regulate neuronal excitability, and their dysfunction causes severe childhood disorders. Here, we identify a structural determinant in the intracellular domains that is responsible for triggering channel activation in response to sodium ion binding. An increase in the size of a particular amino acid renders the channel sodium-insensitive, whilst a decrease in size enables the channel to activate in the absence of sodium. This enhances our understanding of how this subclass of potassium channels respond to changes in the intracellular ionic environment. Furthermore, this may also further our understanding of the basis of human neurological disorders and their treatment.

## Introduction

Na^+^-activated K^+^ channels (K_Na_) open in response to elevation in the cytoplasmic Na^+^ concentration, contributing to hyperpolarization of the membrane potential of neurons and other cell types. K_Na_1.1 and K_Na_1.2 (or SLACK and SLICK) are members of the large-conductance, RCK domain-containing subfamily of K^+^ channels and are activated by intracellular Na^+^ (1). Intracellular Na^+^ is an essential determinant of wild-type (WT) K_Na_1.1 channel opening, and in normal physiology K_Na_1.1 activity is coupled to persistent inward Na^+^ currents, for example through re-opening voltage-gated Na^+^ channels or NALCN channels that increase local Na^+^ concentration above that of the bulk cytosol (2, 3). Additionally, transient currents through AMPA receptors located in the vicinity of K_Na_1.1 have also been implicated as a Na^+^ source for the channel, as part of a negative-feedback loop (4). The importance of these channels is highlighted by the seizure disorders and intellectual disability caused by pathogenic variants in either K_Na_-encoding gene, *KCNT1* or *KCNT2* (5-9). In most cases, pathogenic variants in either gene result in a missense mutation that leads to enhanced K_Na_ channel activity, but loss of function is also found with some *KCNT2* variants (10). Quinidine, bepridil, and clofilium were the first drugs to be identified as K_Na_ channel inhibitors with bithionol, riluzone, loxapine, and niclosamide as activators (11-13). Each of these drugs are non-selective and are unlikely to be of clinical use in K_Na_ disorders. In response to *KCNT1* gain-of-function disorders, several groups have identified novel K_Na_1.1 channel inhibitors (14-16). Our approach exploited K_Na_ channels structures and targeted the pore domain in virtual high throughput screening (14), but it is conceivable that future structure-based screening could instead target the Na^+^-activation mechanism, which could improve specificity. However, the structural basis of how Na^+^ ions interact with K_Na_ channels and how this results in channel activation remained unknown.

Previously, residues located in the rat K_Na_1.1 RCK2 domain were proposed to form a Na^+^ binding site based upon a Na^+^ coordination motif, DXRXXH that is found in Na^+^-activated GIRK channels (17). Mutating D818 and, to a lesser degree, H823 diminished rat K_Na_1.1 Na^+^-activation when the channels were expressed and recorded from patches excised from *Xenopus* ooctyes. In general, Na^+^ sensitivity was shifted to higher concentrations for channels mutated at either site (17). In human K_Na_1.2, mutation of the equivalent aspartate residue, D757, to arginine abolished Na^+^-activation in whole *Xenopus* oocytes, with function rescued by application of the activator niflumic acid (18). The structures of the chicken K_Na_1.1 channel in the closed (zero Na^+^ conditions) and activated (high Na^+^ conditions) conformations were since resolved by cryogenic electron microscopy (cryo-EM) and single particle averaging (19, 20), in which the Na^+^-binding fold proposed by Zhang *et al* (2010) was not evident. The region containing the proposed DXRXXH coordination motif is conserved between chicken and rat K_Na_1.1 but remains as a static loop in both the active and inactive conformations, as will be described below. Since Na^+^ is too small to be identified in the cryo-EM structure, it is not known where in the channel they bind and how this results in increased channel activity.

In studying the cryo-EM structures of the inactive and active chicken K_Na_1.1 channel subunits, we identified a conformational difference between the two that we hypothesized could underlie Na^+^-dependent activation. Furthermore, it appeared to involve the aspartate residue identified by Zhang and others (17) in playing a conformation-stabilizing role. Using site-directed mutagenesis and electrophysiology we identify additional residues in a nearby region that are critical for Na^+^-dependent activation and propose that a twisting of a loop between βN and αQ in RCK2 acts as a molecular switch that underlies K_Na_1.1 activation. Whilst preparing this manuscript, complementary studies on K_Na_1.1 Na^+^-binding and activation were published, involving molecular dynamics simulations and mutagenesis (21) or cryo-EM (22), together with functional characterization. In presenting and discussing our findings we highlight where our interpretation is consistent or conflicts with the conclusions of either of these studies.

## Materials and Methods

### Analysis of protein structures

Structures of chicken K_Na_1.1 in the Na^+^-activated (PDB: 5U70) and Na^+^-free (PDB: 5U76) states (20) were analyzed and figures prepared using UCSF Chimera (23). Amino acid substitutions were introduced *in silico* using the Rotamer function and assessed using the Find Clashes/Contacts function. Molecular surfaces were calculated using the MSMS tool within Chimera (24).

### Molecular biology and transfection

The full-length pcDNA6-K_Na_1.1 mammalian expression plasmid that we described previously (14) was used in these studies. Mutations were designed and introduced by polymerase chain reaction using the New England BioLabs method and confirmed by sequencing (Genewiz, Takeley, U.K.). Due to the large size and high GC content of the insert, mutations were generated in a plasmid containing the SbfI/BsiWI restriction fragment, and then subcloned into the corresponding sites in the pcDNA6-K_Na_1.1 construct. Chinese hamster ovary (CHO) cells were cultured in Dulbecco’s Modified Eagle’s Medium (DMEM) (Gibco, Paisley, UK) supplemented with 10 % (v/v) Fetal Bovine Serum (FBS), 50 U/ml penicillin and 0.05 mg/ml streptomycin and incubated at 37°C in 5% CO_2_. Cells were co-transfected with WT or mutated pcDNA6-K_Na_1.1 together with pEYFP-N1 plasmid using Mirus TransIT-X2 reagent (Geneflow, Lichfield, U.K.). For electrophysiological experiments, cells were plated onto borosilicate glass cover slips and used 2-4 days later.

### Electrophysiology

All chemicals were obtained from Sigma-Aldrich (Gillingham, U.K.) unless stated otherwise. Micropipettes were pulled from thin-walled borosilicate glass (Harvard Apparatus Ltd, Kent, UK), polished, and gave resistances of 1.5 to 2.5 MΩ in the experimental solutions. The bath (extracellular) solution contained, in mM, 140 NaCl, 1 CaCl_2_, 5 KCl, 29 Glucose, 10 HEPES and 1 MgCl_2_, pH 7.4 with NaOH. The 40 mM Na^+^ pipette (intracellular) solution contained, in mM, 130 K-Gluconate, 30 NaCl, 29 Glucose, 5 EGTA and 10 HEPES, pH 7.3 with KOH. To obtain pipette solutions containing Na^+^at 10 mM and 0 mM (nominally Na^+^-free), the NaCl was replaced by equimolar amounts of choline chloride. K_Na_1.1 activators loxapine succinate and niclosamide were prepared as 10 mM stock solutions in DMSO. The final drug concentrations were obtained by diluting the stock solution in the bath solution on the day of the experiment.

Currents were recorded from EYFP-fluorescing cells at room temperature (20 to 22°C) in the whole-cell patch clamp configuration using an EPC10 amplifier (HEKA Electronics, Lambrecht, Germany), with >70 % series resistance compensation (where appropriate), 2.9 kHz low-pass filtering, and 10 kHz digitization. Following the establishment of the whole cell configuration, cells were held at −80 mV and 400 ms voltage pulses from −100 to +80 mV in 10 mV increments were applied. With experiments that examine the effect of pharmacological activation, the 10 mM NaCl pipette solution was used, and the voltage protocol was applied both before and after bath perfusion of 30 μM of either niclosamide or loxapine.

### Data analysis

Samples were not randomized and the experiments were not blinded. Data are presented as mean ± SEM from n number of cells. Statistical analysis was performed using SPSS (IBM Analytics, Portsmouth, U.K.), with the chosen tests indicated in figure legends; *p<*0.05 was considered significant. Representative whole-cell current traces were plotted, and residual capacitance spikes removed in OriginPro. Whole-cell current-voltage relationships were divided by whole-cell capacitance to give current density (pA/pF). Reversal potentials were obtained by fitting the linear part of current-voltage relationships around the reversal potential using linear regression and determining the voltage at the zero current level. Conductance values (*G*) at each voltage (*V*_*m*_) were obtained by dividing current amplitudes (*I*) by the driving force on K^+^ ions, calculated using the reversal potentials (*V*_*rev*_) obtained in individual recordings: *G*=*I*/(*V*_*m*_−*V*_*rev*_). The conductance values were plotted against V_m_ and fitted with a Boltzmann function, *G*=(*G*_*max*_−*G*_*min*_)/(1+e^(*V* −*V*^_*0*.*5*_^)/*k*^)+*G*_*min*_, which gave values for activation midpoint (*V*_*0*.*5*_), *G*_*max*_, *G*_*min*_, and Slope factor (*k*). Data were normalized by dividing by *G*_*max*_ for each experiment. Reported V_0.5_ values were corrected for liquid junction potential error after the experiment. With *k* = *RT/zF*, the valence of the gating charge, *z*, was estimated.

## Results

The aspartate residue found by Zhang and others (17) to be critical for Na^+^ activation in rat K_Na_1.1, D818, is the equivalent to D812 in chicken and D839 in human K_Na_1.1. To assist comparisons between studies of K_Na_1.1 clones from different species the positions of the amino acids detailed in this study in human, rat, and chicken are provided in Table 1. We hereon refer only to the amino acid numbering in human K_Na_1.1. The proposed Na^+^-coordinating motif, based on that in GIRK channels, is conserved between species and forms the loop between βL and αO in the K_Na_1.1 RCK2 domain. Firstly, the Na^+^-binding fold proposed by the molecular modelling and simulations of RCK2 of the rat K_Na_1.1 (17) subunit was not observed in the cryo-EM structures of either the active (PDB: 5U70) or inactive (PDB: 5U76) chicken K_Na_1.1 channels. Secondly there was no discernible difference between the two structures in the positioning of the sidechain of this aspartate or any of the neighboring residues previously proposed to contribute to Na^+^-binding (D839 in Figure 1B). Notably, in the structure of the Na^+^-activated state, a lysine reside in the loop between βN-^α^Q (K885 in Figure 1B) falls within 3.4 Å of the aspartate, sufficiently close to form a salt bridge. In the apo state this loop is not fully resolved, indicating mobility, but the structures suggest a near half-rotation of this loop. Consequentially, the adjacent aspartate in the same βN-αQ loop (D884 in Figure 1B) adopts a position within 4 Å of several acidic or polar residues in the Na^+^-activated state (Figure 1B), which both we and Xu and others (21) termed an “acidic pocket”. Specifically, the aspartate side chain is predicted to contact both E920 and T922 sidechains and the connecting peptide backbone in the βO beta sheet in RCK2 between these two amino acids. The cryo-EM structures therefore indicate that the rotation and stabilization of the βN-αQ loop is a conformational change in RCK2 upon Na^+^ binding.

**Table 1.** Equivalent amino acid positions in K_Na_1.1 channel subunits from the human clone used for electrophysiological experiments in this study and in cryo-EM (22), cryo-EM structures of chicken K_Na_1.1 (20), and rat K_Na_1.1 used in previous studies of Na^+^-activation of K_Na_1.1 (17, 21).

| Human | Chicken | Rat |
| --- | --- | --- |
| E392 | E371 | E373 |
| D839 | D812 | D818 |
| D884 | D857 | D863 |
| K885 | K858 | K864 |
| D898 | D871 | D877 |
| E920 | E893 | E899 |
| T922 | T895 | T901 |

**Figure 1.**
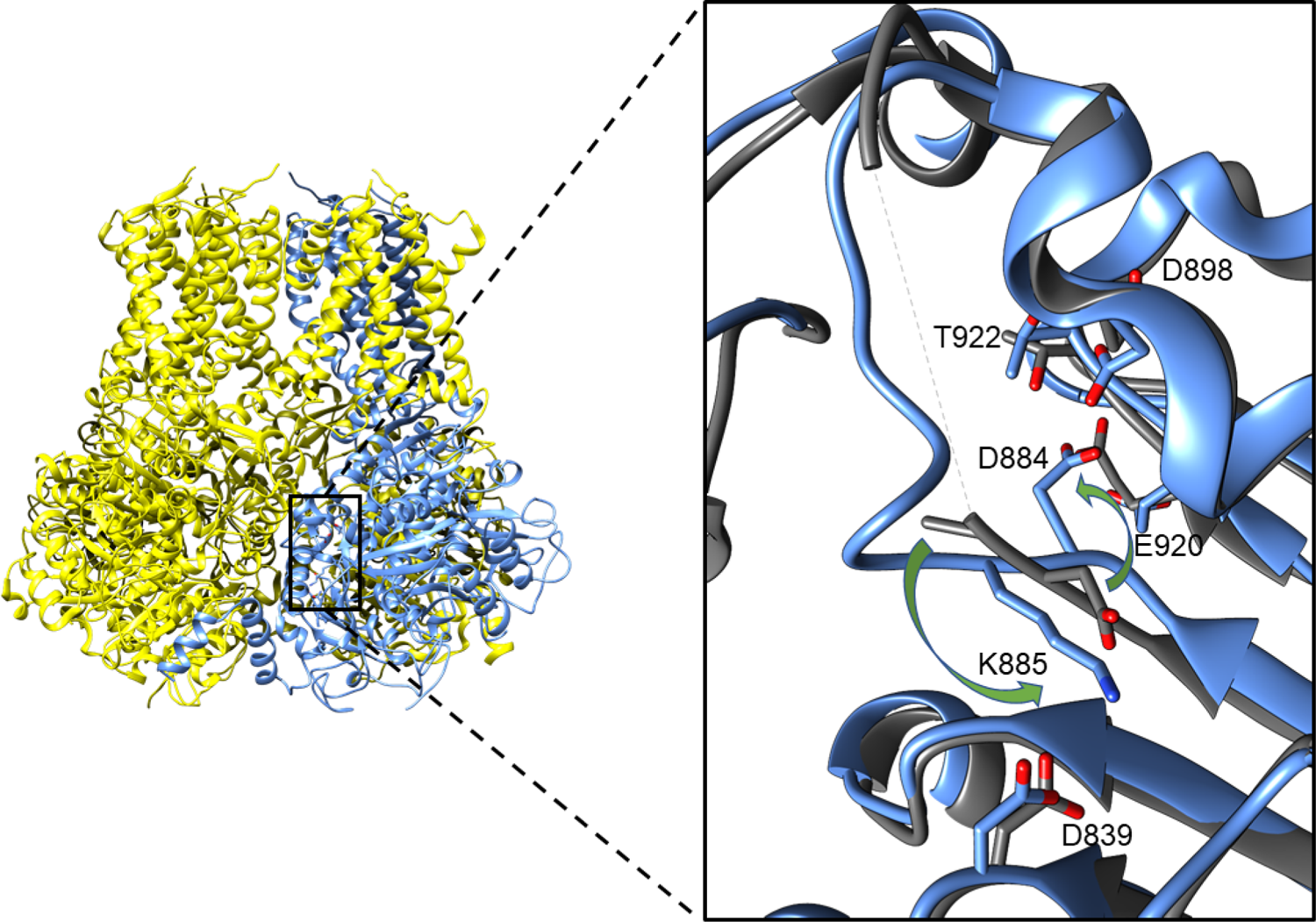
Structural analysis of the Na^+^-binding site proposed by Zhang and others (17) in chicken cryo-EM structures. **A** Structure of chicken K_Na_1.1 in the active conformation (PDB: 5U70) with one subunit in the tetramer colored blue. Each subunit comprises 6 transmembrane helices (upper part of structure), and a re-entrant P-loop between S5 and S6 that forms the K^+^-selective filter. The large intracellular domains contain a pair of RCK domains, RCK1 and RCK2 (lower part of structure). The domain of interest in RCK2 is enclosed by the black box. **B** Enlarged representation of the domain of interest with the active (blue) and inactive (PDB:5U76, dark gray) conformations overlaid. The aspartate identified by Zhang and others (17) is D839 (all human K_Na_1.1 numbering). The twisting of a nearby loop between βN-αQ in RCK2, indicated by the green arrows, with a lysine (K885) close enough to form a salt bridge with the purported Na^+^-binding aspartate (D839) in the Na^+^-activated state. Concurrently, an aspartate (D884) in the same loop adopts a position within 4 Å of several acidic or polar residues. This loop is not resolved in the inactive state (dashed line), suggesting mobility.

### Activation of WT K_Na_1.1 by intracellular Na^+^ and pharmacological activators

Guided by K_Na_1.1 channel structural data, a combination of mutagenesis and whole-cell electrophysiology was used to investigate the involvement of residues in Na^+^-dependent activation of the human K_Na_1.1 channel. In our hands, human K_Na_1.1 runs down within seconds in excised inside-out patches, leaving unitary or no currents. Therefore, to efficiently explore the effects on macroscopic K_Na_1.1 currents, we conducted whole-cell patch clamp recordings from CHO cells expressing WT or mutant K_Na_1.1 using pipette solutions with different concentrations of Na^+^ and using pharmacological activation to bypass Na^+^-activation and confirm the presence of relatively Na^+^-insensitive channels. The anti-psychotic drug loxapine and anti-helminthic drug niclosamide are potent activators of WT K_Na_1.1 with EC_50_ values of 4.4 μM and 2.9 μM, respectively (13). Both drugs also reduce the voltage-dependence of K_Na_1.1 activation, resulting in near-linear current-voltage relationships and increased inward current at voltages negative to the reversal potential (Figure 2). Initially, niclosamide was selected to confirm function expression of “inactive” mutant channels. CHO cells are believed to have little or no endogenous ion conductance (25). However, when 30 μM niclosamide was perfused into the bath a current with density 58.96 ± 9.60 pA/pF at +10 mV was recorded from non-transfected CHO cells, compared to 1.49 ± 0.67 pA/pF at +10 mV prior to its application (n=4 and 5 cells, respectively). Though this current is relatively small in comparison to exogenous K_Na_1.1 current (Figure 2B), this could confound the functional rescue of seemingly inactive K_Na_1.1 channels. The identity of the conductance and charge-carrier evoked by niclosamide is unknown, but experiments ruled out K_Na_1.1, since the current was not inhibited by 10 μM bepridil (not shown). The reversal potential (−73.30 ± 0.85 mV, n=4) would be consistent with a K^+^-selective conductance, but no further experiments were conducted to characterize the current and its ion selectivity. Loxapine had no effect on the membrane conductance of non-transfected CHO cells (Figure 2) and was therefore more suitable for these experiments.

**Figure 2.**
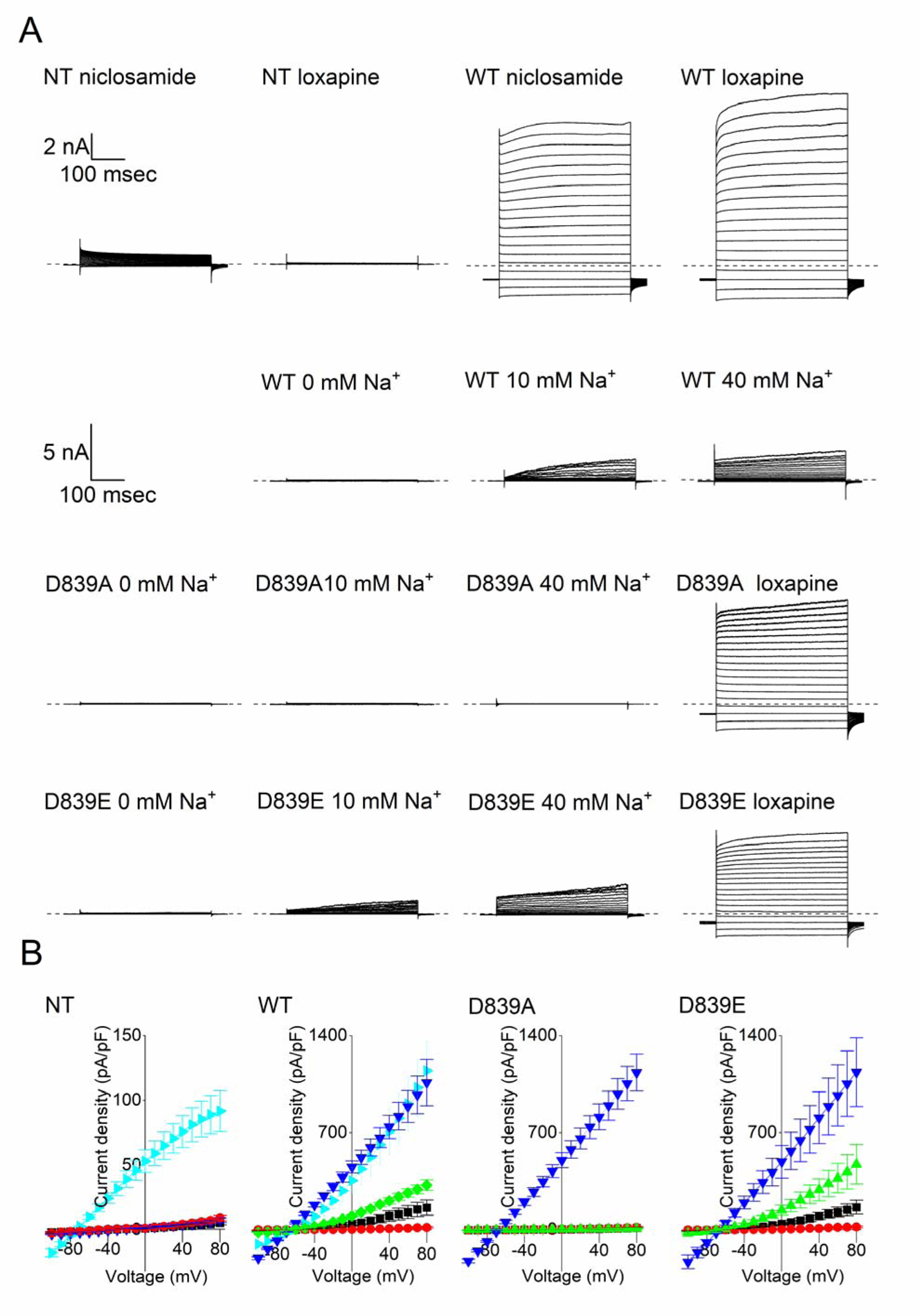
Mutational analysis of the previously proposed Na^+^ sensor. **A** Representative whole cell currents from non-transfected CHO cells (NT) and CHO cells transfected with WT or mutant human K_Na_1.1, as indicated, in response to 400 ms steps from −100 to +80 mV in 10 mV increments from a holding potential of −80 mV. The dashed lines indicate the zero-current level. **B** Mean (± SEM, n=5 to 8 cells) current-voltage relationships for NT control, WT, and mutant K_Na_1.1 channels in the presence of 0 (red 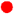), 10 (black 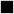) and 40 (green 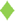 or 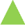) mM intracellular Na^+^, or 10 mM Na^+^ and 30 μM loxapine (dark blue 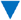). For NT CHO cells and WT K_Na_1.1, 10 mM Na^+^ and 30 μM niclosamide was initially utilized (cyan 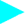) but activated an endogenous CHO cell conductance.

Firstly, to replicate the importance of the previously proposed Na^+^-sensor, D839 was mutated both to glutamate and alanine. The D839E mutation, which lengthened the sidechain without affecting the negative charge, gave currents that were qualitatively no different to WT K_Na_1.1. The current-voltage relationships closely resembled the WT K_Na_1.1 channel with 10 mM and 40 mM intracellular Na^+^, and no currents were recorded in 0 mM intracellular Na^+^ (Figure 2B). Consistent with Zhang and others (17), no currents were recorded in the presence of 0, 10 and 40 mM intracellular Na^+^ from D839A channels. Large, relatively voltage-independent currents were yielded upon perfusion of 30 μM loxapine into the bath solution, confirming the presence of this Na^+^-insensitive mutant K_Na_1.1 channel at the cell membrane (Figure 2).

### Na^+^-activation stabilizes D884 in an acidic pocket

The structural analysis conducted above suggested that Na^+^-activation causes the βN-αQ loop to transition from a highly mobile or disordered state to a conformation that is stabilized by a salt bridge between D839 and K885. To test this idea, K885 was neutralized to alanine. Like with D839A K_Na_1.1, negligible currents were recorded from cells expressing K885A K_Na_1.1 with 0 mM and 10 mM intracellular Na^+^ (Figure 3). A small current was recorded from K885A K_Na_1.1 when intracellular Na^+^ was elevated to 40 mM, suggesting a substantial decrease in Na^+^-sensitivity of the channel. Functional expression of K885A K_Na_1.1 channels was again confirmed by addition of 30 μM loxapine, which evoked large whole cell currents comparable with those we observed earlier in the study.

**Figure 3.**
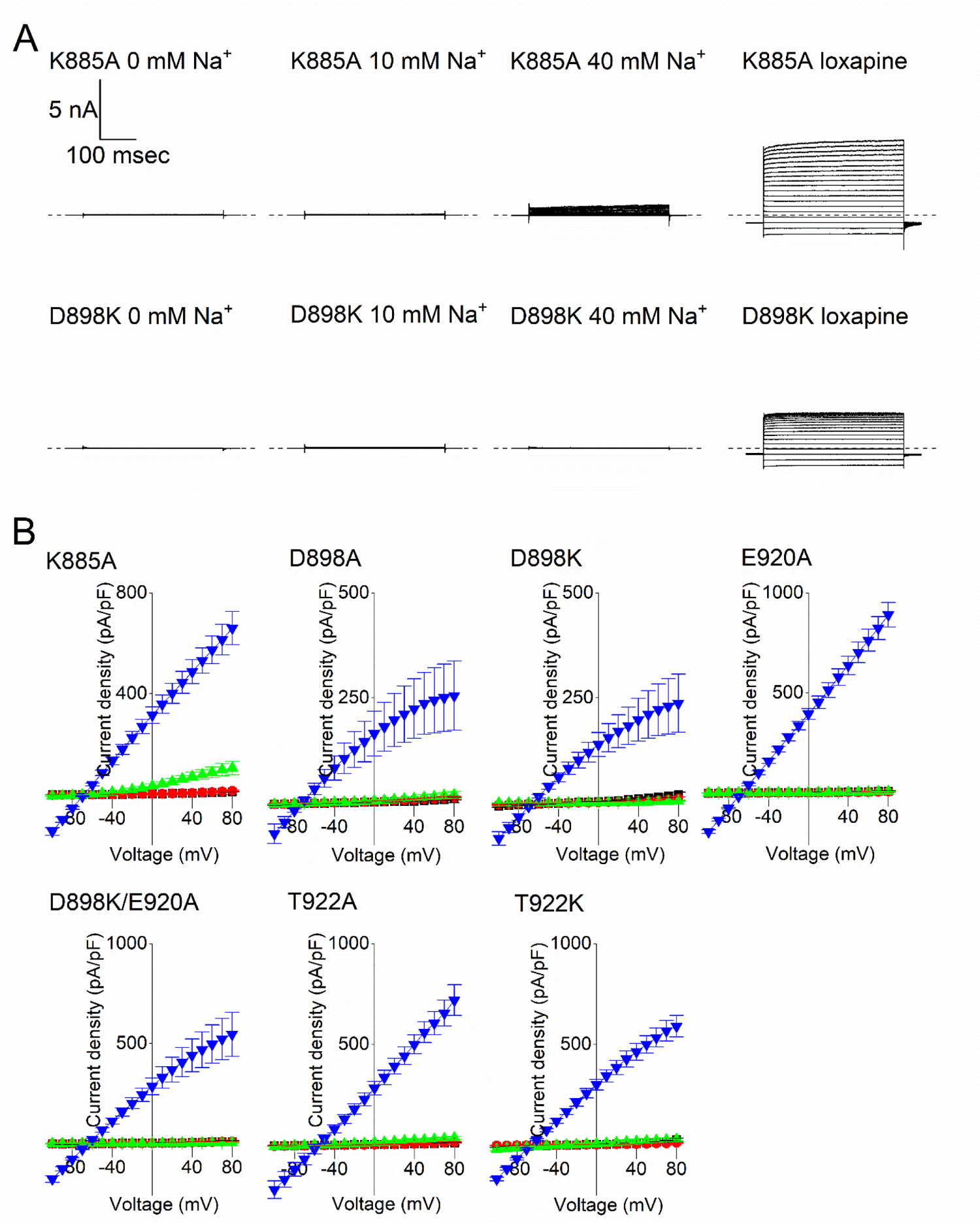
Disruption of Na^+^ activation through loss of the D839/K885 salt bridge and acidic pocket residues. **A** Representative whole cell K885A and D898K K_Na_1.1 currents in each condition, as indicated, in response to 400 ms steps from −100 to +80 mV in 10 mV increments, from a holding potential of −80 mV. The dashed lines indicate the zero-current level. **B** Mean (± SEM, n=5 to 8 cells) current-voltage relationships for mutated K_Na_1.1 channels in the presence of 0 (red 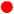), 10 (black 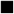) and 40 (green 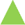) mM intracellular Na^+^, or 10 mM Na^+^ and 30 μM loxapine (dark blue 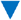).

We then questioned if the concurrent positioning of D884 in the vicinity of other negatively charged (D898 and E920) or polar (T922) sidechains was significant. Mutation of each of these to alanine similarly disrupted Na^+^ activation with no significant currents recorded with 0, 10, or 40 mM intracellular Na^+^. Functional rescue was achieved with each of these mutant K_Na_1.1 channels following application of 30 μM loxapine. We initially considered whether this acidic pocket might form a site for Na^+^ to occupy and activate the channel, co-ordinated by these several negatively charged and polar residues. To see if we could rescue one of these mutations, E920A, or attract D884 into this site and increase activity, D898 and T922 were mutated to the positively charged lysine. However, with each of the D898K, E920A/D898K, and T922K K_Na_1.1 channels the effect was same as the alanine substitutions in this pocket (Figure 3). Again, no currents were recorded with 0, 10, or 40 mM intracellular Na^+^ and functional rescue was achieved with 30 μM loxapine application. We found that that mutation of D898 to either alanine or lysine appeared to result in a weak inward rectification in loxapine-evoked current-voltage relationships (Figure 3B).

### The size of amino acid substitutions of D884 in the βN-αQ loop controls the activation state of K_Na_1.1 channels

We returned to investigate the significance of D884 and its negative charge, which occupies the acidic pocket upon Na^+^ activation, by mutating to alanine. Unexpectedly, this resulted in a large increase in channel activity and apparent loss of Na^+^-dependence. The voltage-dependent currents recorded from CHO cells expressing D884A K_Na_1.1 with 0, 10 or 40 mM intracellular Na^+^ were all similar (Figure 4A). Perfusion of 30 μM loxapine to cells had little effect on the current amplitude apart from an apparent loss of voltage-dependence, as indicated by a straightening of the current-voltage relationship (Figure 4B). Mutation of the same residue to valine, which has a longer sidechain by just one carbon, had similar effects (Figure 4). Activation midpoints derived from conductance-voltage relationships fitted with a Boltzmann equation for D884A and D884V in 10 mM Na^+^ were significantly hyperpolarized compared to WT K_Na_1.1 (Table 2). With 40 mM intracellular Na^+^, the activation midpoints for D884A and D884V K_Na_1.1 were not significantly different from WT K_Na_1.1 under the same conditions. Whilst WT K_Na_1.1 currents were negligible with 0 mM intracellular Na^+^, the activation midpoints for D884A and D884V with 0 mM Na^+^ were not significantly different to those obtained with 10 and 40 mM Na^+^ (Table 2).

**Table 2:**
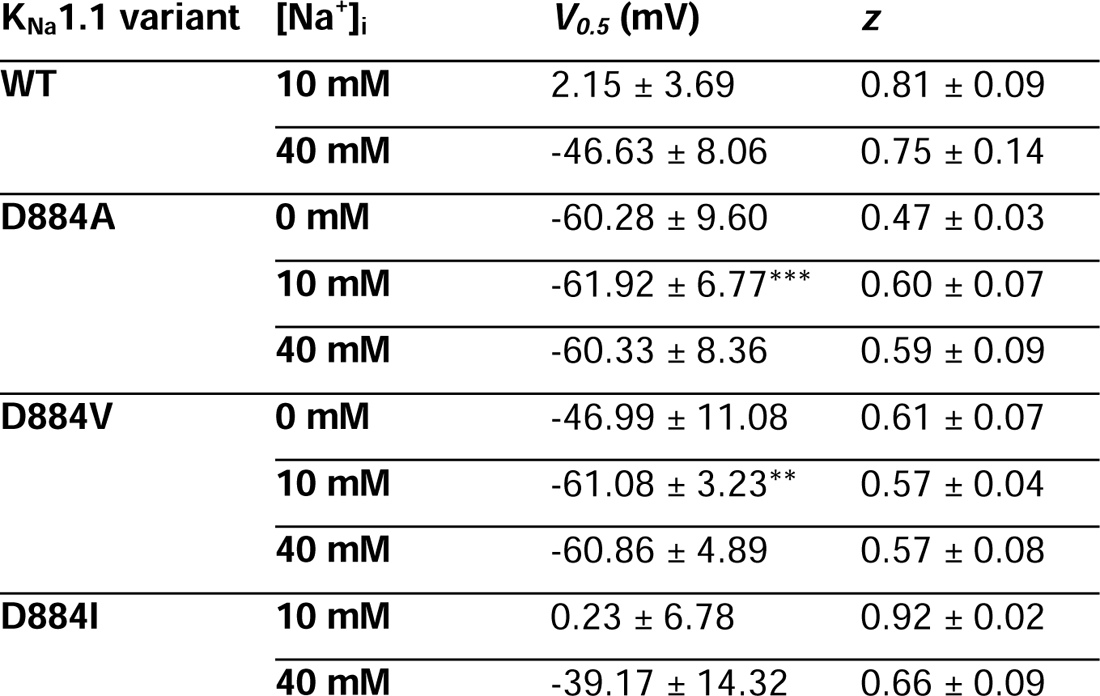
Parameters derived from Boltzmann fit of WT and D884 mutant K_Na_1.1 channels with 0, 10, and 40 mM intracellular Na^+^. Data are presented as mean ± SEM (n=5 to 9 cells). *V*_*0*.*5*_ is the half-maximal activation voltage, ^\*\**p*<0.005, ***^*p*<0.0005 compared to WT V_0.5_ with 10 mM intracellular Na^+^ (independent one-way ANOVA with Tukey’s post-hoc test). *z* was derived from the slope of the Boltzmann curve, *RT*/*zF*.

**Figure 4.**
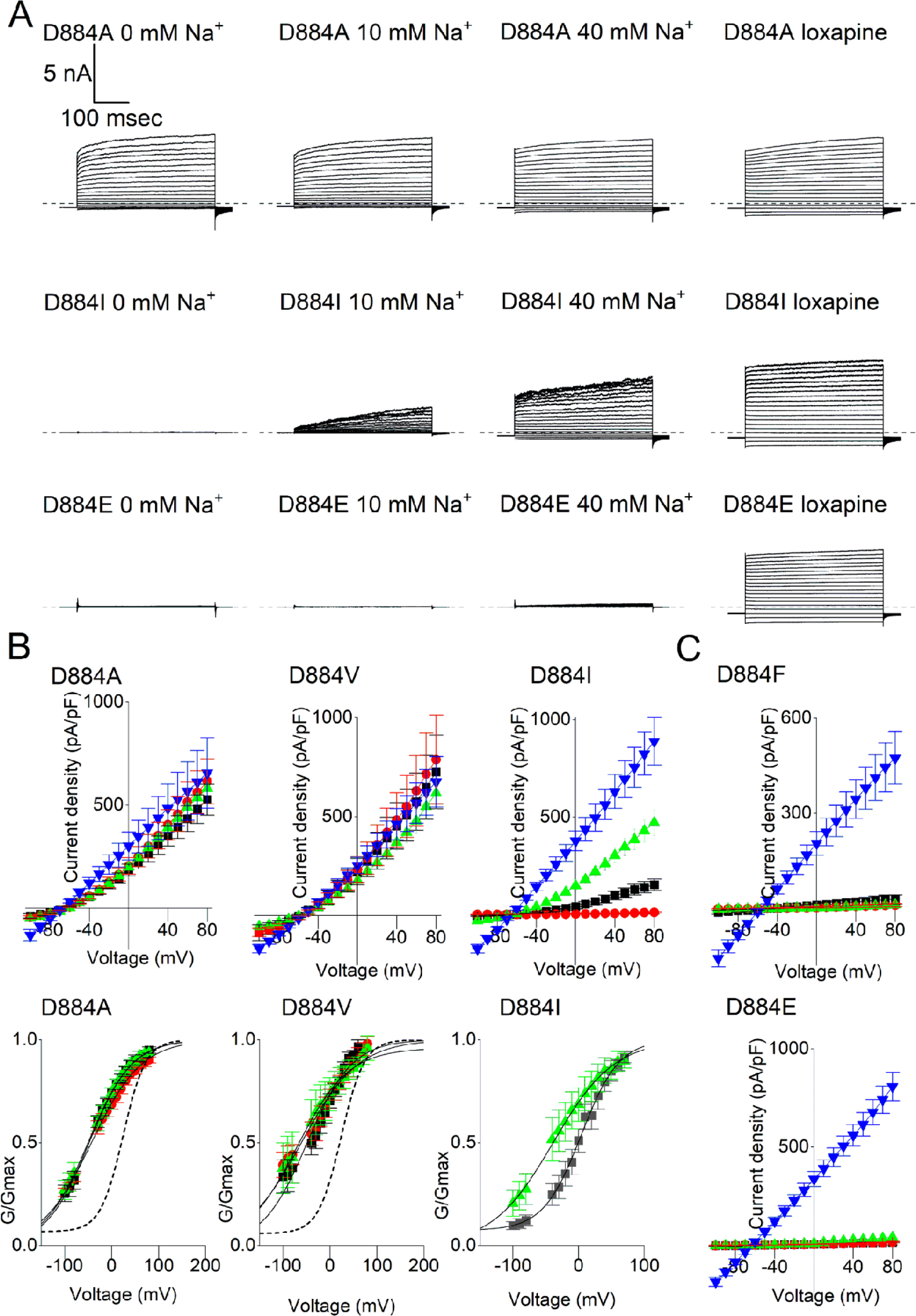
Role of the K_Na_1.1 amino acid size at position 884 in controlling channel activation. **A** Representative D884A, D884I, and D884E K_Na_1.1 whole cell currents in response to 400 ms steps from −100 to +80 mV in 10 mV increments, from a holding potential of −80 mV. The dashed lines indicate the zero-current level. **B** Mean ± SEM (n=5 to 8 cells) current-voltage and conductance-voltage relationships for D884A, D884V, and D884I K_Na_1.1 with 0 (red 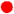), 10 (black 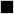) and 40 (green 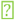) mM intracellular Na^+^, or 10 mM Na^+^ and 30 μM loxapine (dark blue 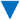). The mean conductance-voltage relationship for WT K_Na_1.1 recorded with 10 mM intracellular Na^+^ is indicated by a dotted line for comparison. **C** Mean ± SEM current-voltage relationships for D884F and D884E K_Na_1.1 with 0 (red 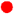), 10 (black 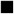) and 40 (green 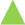) mM intracellular Na^+^, or 10 mM Na^+^ and 30 μM loxapine (dark blue 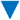).

Upon increasing the size of the hydrophobic sidechain further by mutating D884 to isoleucine and phenylalanine, different effects were observed. D884I K_Na_1.1 currents were similar to those obtained from WT K_Na_1.1 with each intracellular Na^+^ concentration tested (Figure 4B, Table 2). In contrast, no discernible current could be recorded from D884F K_Na_1.1 with any of the Na^+^ concentrations tested up to 40 mM, though function was rescued by loxapine application (Figure 4B). It appeared that it was the size of the sidechain at this position that was critical to determining K_Na_1.1 channel function. Finally, we mutated D884 to glutamate, which retained the negative charge but increased the sidechain by one carbon. The D884E mutation in K_Na_1.1 had an effect similar to D884F, with negligible currents with each of 0, 10, and 40 mM intracellular Na^+^, but large whole-cell currents were recorded following the application of loxapine (Figure 4C).

These results indicated that it was the size and not the charge of the side chain at amino acid position 884 that determined whether the K_Na_1.1 channel remained Na^+^-dependent or became locked in either the inactive or Na^+^-activated state under these conditions. We returned to the structure of the Na^+^-activated K_Na_1.1 channel and modeled the D884 mutations *in silico*. Whilst aspartate and isoleucine at position 884 could be closely accommodated in the surrounding pocket, together with the smaller alanine and valine sidechains with some leeway, substituting the larger glutamate and phenylalanine sidechains resulted in steric clashes with T922 (Figure 5). D884 was mutated to asparagine in rat K_Na_1.1 by Xu and others (21), resulting in Na^+^-insensitivity, and we found that this mutation also resulted in a steric clash *in silico* (not shown).

**Figure 5.**
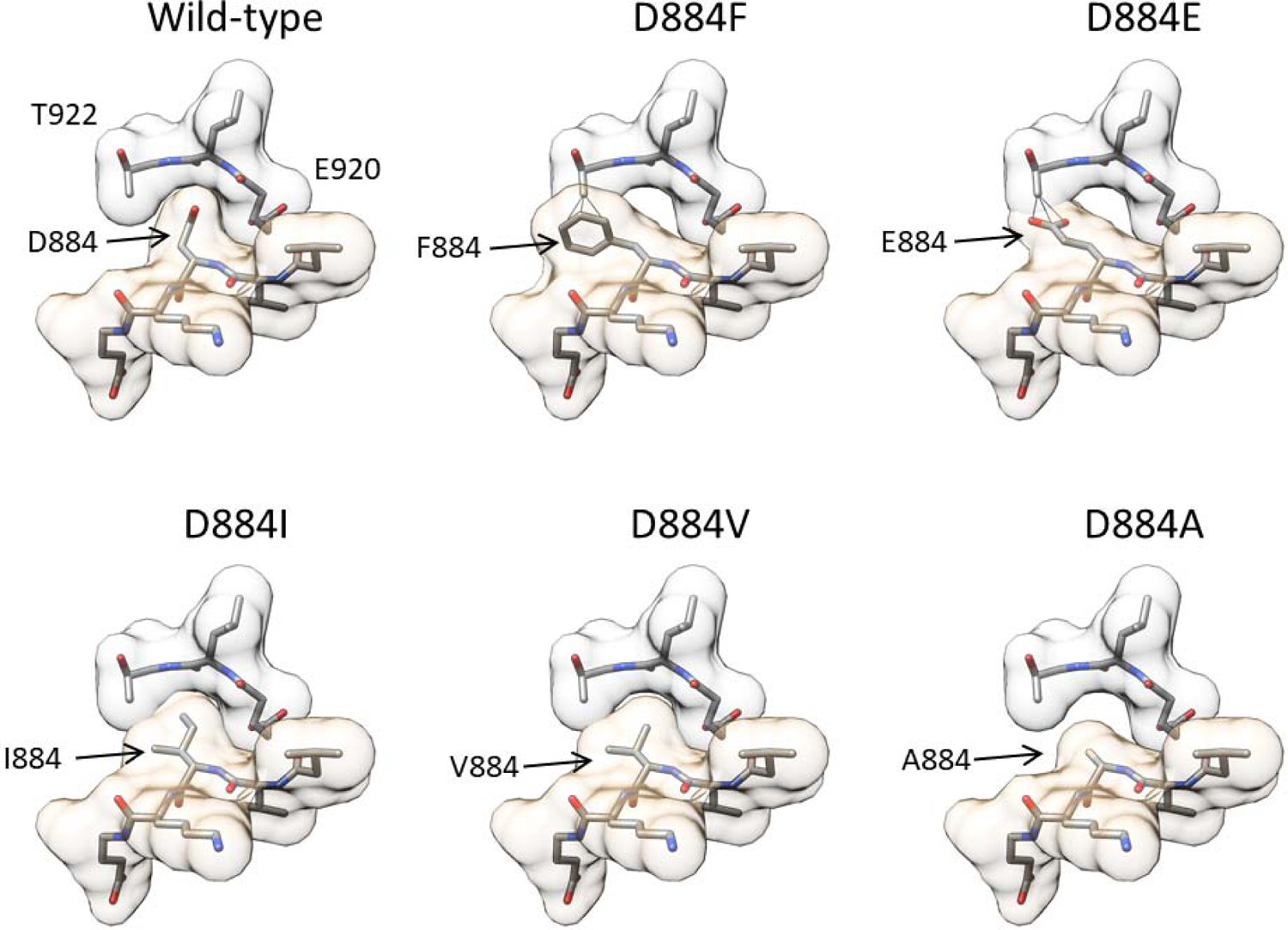
Molecular modelling of D884 mutations. Models were generated from the structure of chicken K_Na_1.1 in the Na^+^-activated state (PDB: 5U70), here with the numbering of conserved residues in human K_Na_1.1. In each model the pocket formed by ^920^ELT^922^ in βO is shown with pale gray space fill, with ^882^VVDKE^886^ in the βN-αQ loop with pale gold space fill below. With the D884F and D884E substitutions, predicted atomic clashes between the substituted sidechains and T922 are indicated by solid black lines. No clashes were predicted in the other models.

## Discussion

Our data suggest that the βN-αQ loop in RCK2 of K_Na_1.1 acts as a molecular switch, controlling transitions between the Na^+^-activated and Na^+^-free inactive states. In the activated state, this loop is stabilized by a salt bridge between K885 and D839. This explains why previous mutations of this aspartate, D812 in rat K_Na_1.1, by Zhang and others (17) resulted in a loss of Na^+^-sensitivity, leading to their proposal that this formed a Na^+^-binding site. Concurrently, the ability of D884 to occupy a pocket that includes D898, E920, and T922 in the activated state seems to be important for activation. Increasing the size of this sidechain prevented K_Na_1.1 channel activation by 40 mM intracellular Na^+^ but decreasing the size of the sidechain caused the channel to adopt a constitutively activated, but voltage-gated, Na^+^-independent state. This would imply that the conformation adopted by this pocket in the absence of Na^+^ prevents D884 and the βN-αQ loop from adopting the stabilized state, but can accommodate alanine and valine sidechains in place of D884. This domain is highly conserved between K_Na_1.1 and K_Na_1.2, including each of the residues described here, indicating a common mechanism.

This acidic pocket together with D884 were recently proposed by both Xu and others (21) and Zhang and others (22) to be a potential cation binding site. Mutating the equivalent of D884 in rat K_Na_1.1 to asparagine rendered the channel non-functional at intracellular Na^+^ concentrations to up to 2 M (21). Consequently, and supported by molecular dynamics simulations, this aspartate was proposed as one of two Na^+^-binding sites in each subunit. In contrast, we found that the charge of this residue is not critical to K_Na_1.1 channel function, since D884I K_Na_1.1 channels behaved similarly to WT K_Na_1.1. *In silico* mutagenesis of this aspartate in the chicken K_Na_1.1 structure to asparagine caused a steric clash with acidic pocket residues, similar to D884E and D884F, which substitutions prevented Na^+^-activation in our experiments. This explains the lack of activity with the asparagine mutation observed by Xu and others (21). D884 is part of cation-binding “site 2” identified by Zhang and others (22), who also studied the D884A K_Na_1.1 mutation. Using intracellular Na^+^ concentrations upwards from 100 mM, they found a four-fold decrease in the Na^+^ EC_50_. Since we recorded activated K_Na_1.1 channel currents in the absence of Na^+^, this may represent residual Na^+^-dependence from an alternative Na^+^ binding site with these higher concentrations.

We are unable to propose a specific Na^+^-binding site from our investigation, but mutagenesis of acidic pocket residues D898, E920 and T922 that also disrupted Na^+^ activation would support the view that Na^+^ binds in this region. These residues may have a role in co-ordinating Na^+^ ions, consistent with E920 having been identified by Xu and others (21) as a Na^+^ co-ordinating sidechain. Mutation of one of these residues, D898, to either lysine or alanine resulted in inward rectification of the K_Na_1.1 current-voltage relationship when activated by loxapine. It was noted in the structure of the chicken K_Na_1.1 channel that the ring of RCK domains forms a “funnel” that narrows as it approaches the pore-forming region and has a largely electronegative surface (19). D898 contributes to this highly negatively charged surface which may contribute to ion conduction. The negatively charged residues are thought to attract K^+^ ions and contribute to the relatively high K^+^ conductance of this and other member of the channel subfamily, and mutation of D898 may have an effect of outward K^+^ currents, leading to the slight inward rectification observed.

The rotation and adoption of a stable alpha helix by the βN-αQ loop in RCK2 was also highlighted in the recently published structure of the human K_Na_1.1 in the activated state, but disordered in the Na^+^-free state (22). The authors argue that this conformational change influences interactions between RCK2 and RCK1 domains, resulting in the overall expansion that opens the channel gate. In conclusion, this structural and our functional data point to this loop as the key regulator of K_Na_ channel activation by Na^+^. The constitutive activity with the D884A and D884V mutant K_Na_1.1 in the absence of Na^+^ provide further clues to how this channel is regulated. Given that these mutant channels, with the smaller side chain, are able to activate without Na^+^, this indicates that the WT channel is primed for activation but requires a small structural change caused by Na^+^ binding to enable the βN-αQ loop to adopt the activated state. It is this inactive, but primed state that may be disrupted by inherited mutations that result in K_Na_1.1 gain of function.

## Author contributions

BAC, JDL, ACK, and SPM designed the study. BAC and JDL performed research. BAC, JDL, ACK, SPM, and NP analyzed and interpreted data. BAC and JDL drafted the manuscript BAC, JDL, ACK, SPM, and NP edited and approved the manuscript.

## Declaration of Interests

The authors declare no competing interests.

## Acknowledgments

Supported by a BBSRC-CASE PhD studentship in conjunction with Autifony Therapeutics Ltd awarded to B.A.C. (BB/M011151/1). For the purpose of open access, the authors have applied a Creative Commons Attribution (CC BY) license to any Author Accepted Manuscript version arising from this submission.

## Notes

### Competing Interest Statement

The authors have declared no competing interest.

### Summary of Updates

Replaced symbols in some figures that failed to convert to PDF. Figure 5 amino acid numbering corrected.

